# Pangenome evaluation of gene essentiality in *Streptococcus pyogenes*

**DOI:** 10.1101/2023.08.29.555273

**Authors:** Magnus G. Jespersen, Andrew J. Hayes, Steven Y. C. Tong, Mark R. Davies

## Abstract

Populations of bacterial pathogens are made of strains that often have variable gene content, termed the pangenome. Variations in the genetic makeup of a single strain can alter bacterial physiology and fitness in response to different environmental stimuli. To define biologically relevant genes within a genome, genome-wide knockout transposon mutant libraries have been used to identify genes essential for survival or virulence in a particular strain. Such phenotypic studies have been applied in four different genotypes of the major human pathogen *Streptococcus pyogenes*, yet challenges exist in comparing results across studies conducted in different genetic backgrounds and conditions. To advance genotype-phenotype inferences within a population genomic framework of 250 *S. pyogenes* reference genomes, we systematically re-analysed publicly available transposon sequencing datasets from *S. pyogenes* using a transposon sequencing specific analysis pipeline, Transit. Across 4 genetic backgrounds and 9 phenotypic conditions, 311 genes were highly essential for survival, corresponding to ∼22% of the core genome. Among the 311 genes, functions related to information storage, and processing were overrepresented. Genes associated with cellular processing and signalling were of significantly higher essentiality under *in vivo* conditions (animal models with differing disease manifestation and site of colonisation) compared to *in vitro* (varying types of culture media). Finally, essential operons across *S. pyogenes* genotypes were defined, with an increased number of essential operons detected under *in vivo* conditions. This study provides an extendible database to which new studies can be added, and a searchable html-based resource to direct future investigations into *S. pyogenes* population biology.

**Importance:** *Streptococcus pyogenes* is a human adapted pathogen occupying restricted ecological niches. Understanding essentiality of genes across different strains and experimental conditions is important to direct research questions and efforts to prevent the large burden of disease caused by *S. pyogenes*. To this end we systematically reanalysed transposon sequencing studies in *S. pyogenes* using transposon sequencing specific methods, integrating them into an extendible meta-analysis framework. This provides a repository of gene essentiality in *S. pyogenes* for the community to guide future phenotypic studies.

## Introduction

*Streptococcus pyogenes* is a human adapted pathogen estimated to cause more than 517,000 deaths and more than 720 million cases of superficial infections such as pyoderma or pharyngitis each year (1). Development of vaccines to prevent *S. pyogenes* is an area of active research, with no product yet commercialised, despite clear health and economic incentives (2–6). As a result of the high burden of disease, research to understand the fundamental biology of this pathogen is required to define which gene(s) are central to the survival of the pathogen irrespective of the ecological niche occupied or clinical presentation associated with *S. pyogenes*. This knowledge can be used to identify new drug targets or better understand the vast amounts of data generated using high-throughput methods, such as transcriptomics and other -omics techniques.

Transposon sequencing is one high-throughput method that has been successfully used to probe gene essentiality in bacteria. While multiple versions of this method have been developed (7–12), common is the use of modified transposable elements (transposons), which insert semi-randomly into the genome. If a transposon inserts in or close to a gene, this is likely to cause a gene knock-out. Through this approach, a mutant library is generated with each bacterial cell having a transposon inserted ‘randomly’ in its genome, which can then be screened using high-throughput sequencing methods for regions of increased or decreased transposon insertion rate. The rate of transposon insertion indicates essentiality of a specific sequence under a given condition and is detected by increased or decreased mutant survival rates (13). In *S. pyogenes,* transposon sequencing has been used to examine gene essentiality under different *in vivo* and *in vitro* conditions (14–22). These studies have highlighted important genes related to survival, colonization, and infection in varying environments and conditions including human challenge and murine skin or soft tissue infections (18, 23).

One of the limitations of transposon sequencing has been an inability to systematically compare between studies where strain type, conditions, and informatic processing is not uniform. An additional complexity is the fact that *S. pyogenes* is a genetically diverse organism and total gene content varies across different strains (2). Recent transposon studies have focused on producing collections of transposon mutant libraries and analysed these to elucidate pan-genome essentiality where interactions with or between accessory genes can lead to essentiality of one or more genes (24–26). In *S. pyogenes* only few transposon sequencing studies have been put into a co-analysis to increase understanding across datasets. When studies are compared, results often originate from different analysis pipelines possibly introducing additional biases. Here we conducted a systematic re-analysis of transposon sequencing studies from *S. pyogenes*, with methods specifically developed for transposon sequencing (27, 28), and subsequent weighting of each gene in terms of its essentiality. These findings serve as an extendible database for gene essentiality, provided in a html-based format for easy search, to guide future research and to better understand the biology of *S. pyogenes* infections.

## Results

A total of 9 transposon sequencing studies have been conducted in *S. pyogenes* (14–22). Of these, 5 studies representing 9 datasets were analysed in this study after applying the following exclusion criteria: Microarray-based (14); raw reads were not publicly available (17, 21); condition-restricted preventing comparability (20, 22); and low saturation of transposon insertion (15). These datasets included represented four *S. pyogenes* strains of three *emm* types: two *emm*1 strains (5448 and MGAS2221), one *emm* 28 (MGAS27961), and one *emm*49 (NZ131). The datasets covered *in vitro* (n = 4) and *in vivo* (n = 4) growth conditions with *emm*1 and *emm*28 tested across both (Supplementary Figure 1). The *emm*49 strain was only tested *in vitro*. Finally, one dataset of *emm1* consisted of an *in vitro* Zn^2+^ challenge, which was not included in *in vitro* essentiality calculations due to the added stressor. *In vitro* and *in vivo* conditions covered multiple different media types and sites of infection in different model organisms (Table 1). To unify separate datasets we applied optimal trimming, alignment and count of insertions with the tool Transit (27, 28) taking Transposon type into account. This method allowed for systematic assessment of gene essentiality across experimental discrepancies (refer to materials and methods).

**Table 1.**
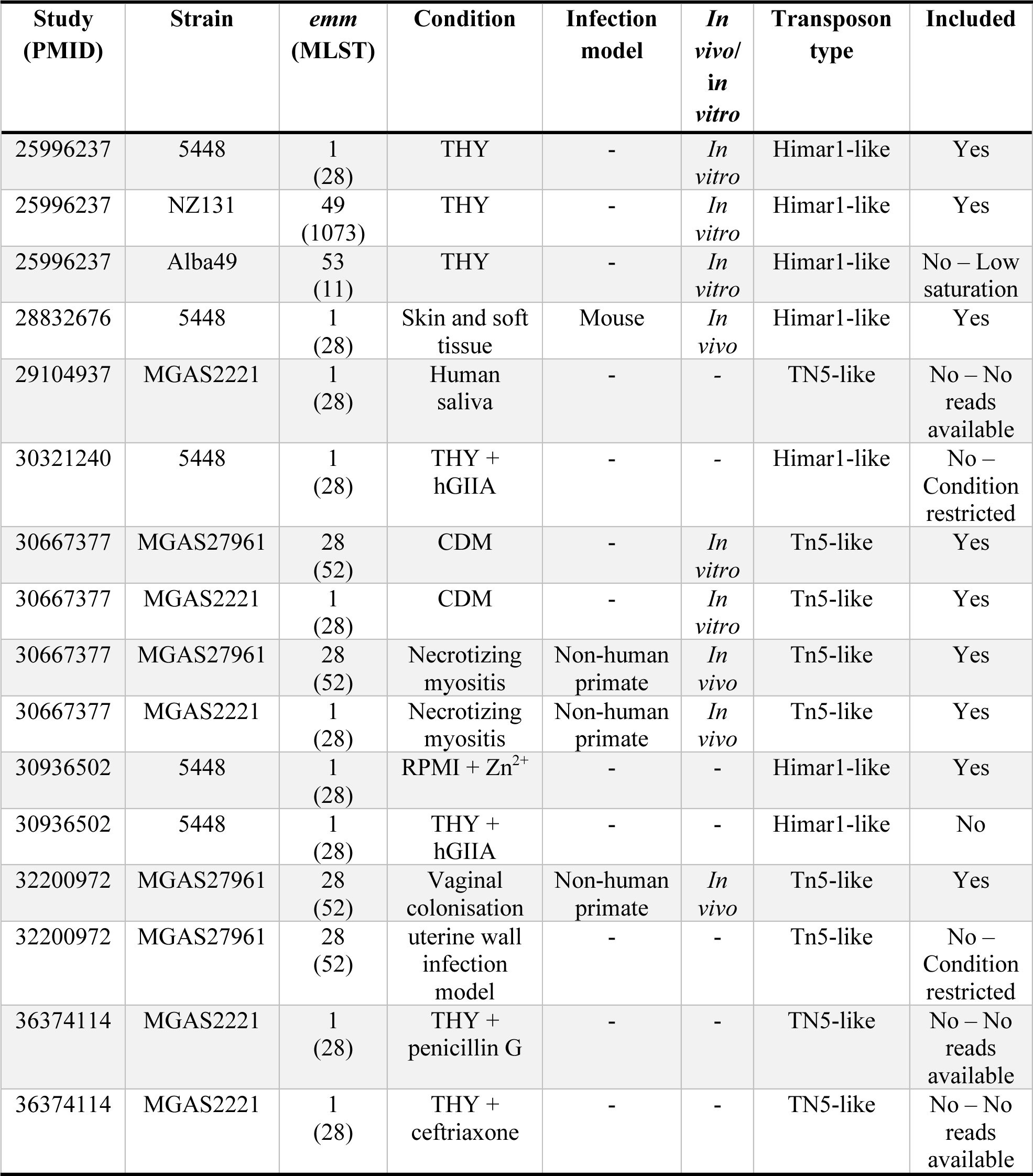
Transposon sequencing studies of *Streptococcus pyogenes*. General overview of studies utilizing transposon sequencing to examine *S. pyogenes*. hGIIA: human group IIA secreted phospholipase A2, THY: Todd-Hewitt medium supplemented with yeast extract. Exclusion criteria: Micro array based; No publicly available raw reads; condition-restricted preventing comparability; and low saturation of transposon insertion.

### S. pyogenes reference pangenome

As existing transposon sequencing studies were performed in different genetic backgrounds, a pangenome approach using all publicly available complete reference genomes was chosen to identify gene orthologs. The rationale for incorporating all available reference genomes was to increase broader utility of the dataset by extrapolating gene essentiality across reference genomes and additional strains of *S. pyogenes*, and to have a framework to observe potential lineage adaptations to variable genes across the population (accessory genes). An *S. pyogenes* ‘reference’ pangenome was constructed using Panaroo (29) from 249 publicly available reference genomes (Supplementary file 1) representing 103 different *emm* types(29). Functional annotation of the pangenome was obtained using EggNOG-mapper and the EggNOG database (30, 31). Of the 4,178 genes identified in the ‘reference’ pangenome, 1,390 (1,390/4,178 - 33.3%) genes were identified as core genes (encoded in ≥99% of isolates (≥246 genomes) (Figure 1A)). 1,999 (1,999/4,178 - 47.8%) genes appeared in at least one of the four strains used in transposon sequencing studies. 2,788 genes were identified as accessory (Figure 1A). Annotations from Eggnog were used to assign Clusters of Orthologous Genes (COG) categories which were collated into major categories based on related function, to simplify interpretation (Supplementary Table 1, Supplementary html 1). Both core and accessory genes were predicted to encode products in each of the major COG categories (Figure 1A).

**Figure 1.**
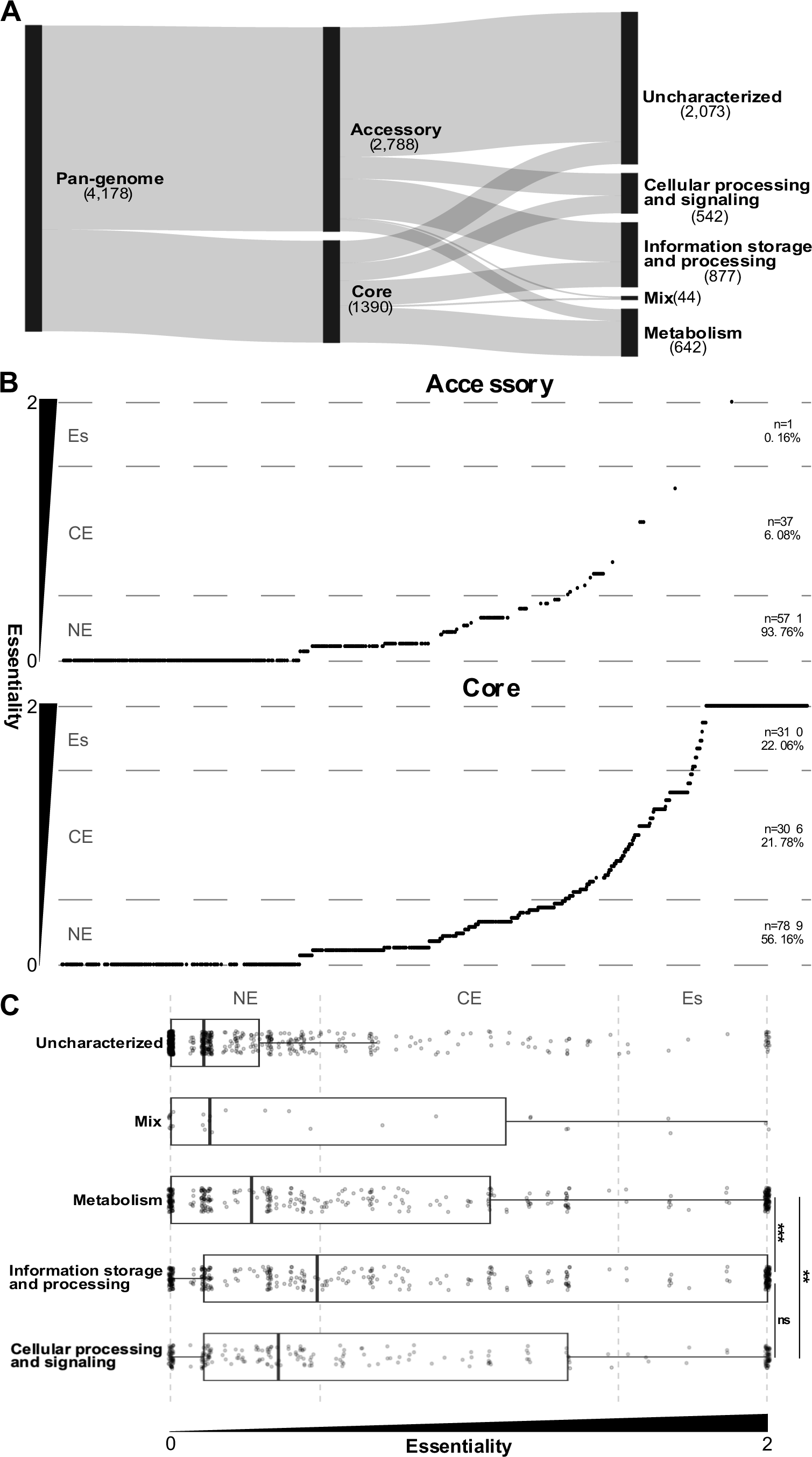
Gene essentiality and functional classification across a Streptococcus pyogenes reference pangenome. **A**: Sankey flow diagram from left to right illustrating how genes from the pangenome of 249 complete genomes, are distributed among core and accessory categories, and different functional categories based on Clusters of Orthologous Gene categories. **B**: Distributions of gene essentiality scores for core and accessory genes, ordered vertically by essentiality score. Vertically, essentiality score increases from the bottom to the top and horizontal grey dashed lines indicate borders between quantitative categories of essentiality (non-essential (NE), conditionally-essential (CE), essential (Es)), as well as the minimum and maximum essentiality values indicated by the bottom and top lines, respectively. **C**: Gene essentiality scores based on functional attributes, where each point represents a single gene, and the box and whisker plots indicating the overall distribution. The essentiality score increases from left to right. Results of statistical comparisons (Dunn’s test) are indicated by lines between compared categories. n.s.: not significant, **: adjusted p-value < 0.01, ***: adjusted p-value < 0.0001.

### Importance of the core genome and unveiled strain adaptations

To identify essential genes across the *S. pyogenes* pangenome, a gene essentiality score was determined. This score is a semi-quantitative measure of essentiality across datasets calculated for a specific gene or collection of genes. Essentiality scores range from 0 (non-essential) to 2 (highly essential). Due to the skew of strains used in publicly available datasets, *emm*1 (n = 4), *emm*28 (n = 4), and *emm*49 (n = 1), the essentiality score for a gene was calculated as the mean of lineage (*emm*) specific essentiality scores. This method aimed to find a consensus essentiality score within each lineage, and then weight these equally to avoid lineage bias. For the 2000 genes represented across the 9 transposon mutagenesis studies, core genes had a significantly larger essentiality score compared to accessory genes (Mann-Whitney U test: p<2.2·10^-16^) (Figure 1B). The median essentiality score for core genes was 0.42 (Q1 = 0.11; Q3 = 1.33) meaning that a little over half of the core genome is seemingly of low essentiality (essentiality score < 0.5, n = 792, 792/1390, 56%), with less than a quarter being of high essentiality (essentiality score > 1.5, n = 309, 309/1390, 22%). For accessory genes the median was 0.00 (Q1 = 0.00; Q3 = 0.133)

### Enriched functional categories and poorly characterised genes

Dividing genes into their major COG categories revealed that in general cellular processing, and signalling (CPS), information storage and processing (ISP), and metabolism were of greater essentiality compared to genes of mixed categories and uncharacterized genes (Figure 1C). Comparing distributions of gene essentiality score for major COG categories revealed metabolism (Q1 = 0.00; median = 0.27; Q3 = 1.07) had a significantly lower essentiality score compared to CPS (Q1 = 0.11; median = 0.36; Q3 = 1.33, Dunn’s post-hoc test: adjusted p=0.008) and ISP (Q1 = 0.11; median = 0.49; Q3 = 2.00, Dunn’s post-hoc test: adjusted p=4.4·10^-6^)(Figure 1C). There was no statistical difference between CPS and ISP (Dunn’s post-hoc test: adjusted p=0.083) (Figure 1C), suggesting that more genes within the functional categories related to CPS and ISP are essential and knockouts of these functions are lethal. One example of a set of highly essential core genes with annotated functions are the fatty acid biosynthesis (*fab*) genes. These 13 genes, organised into three operons (o_0583, o_0584, and o_0585 in S119)(32, 33), are highly essential across all conditions. The *fab* genes have previously been linked to virulence (34, 35). The results presented here and in previous studies stress *fabT* and the remaining *fab* genes as potential targets for narrow range therapeutics (32), highlighting how increased knowledge about cross lineage fundamental functions of *S. pyogenes* can direct discovery and research into therapeutic targets.

In contrast to functionally characterized essential genes, others remain functionally uncharacterised in *S. pyogenes* but none-the-less have high essentiality scores. Some of these genes have been studied in other streptococcal species, while others have no annotated function. Genes with poor annotation and large essentiality scores in this dataset can guide efforts to study genes essential for *S. pyogenes* biology. An example of such a gene is *glpF2* (also called *Aagp*) (SF370 tag: SPY_RS07685/Spy1854). *glpF2* has been identified as an atypical aquaglyceroporin facilitating transport of hydrogen peroxide and likely glycerol (36). In *Streptococcus suis glpF2* was identified as important during infection of mice using a knockout strain, and in pigs by transposon mutagenesis (36, 37). Additionally, a *glpF2* knockout displayed a decreased lag phase duration during growth at 42°C in rich medium (37). Apart from *speA, glpF2* is notably found as having an increased transcript abundance level when comparing multiple M1_Global_ and M1_UK_ strains, prompting the need for a better understanding of *glpF2* and its potential involvement in virulence, survival, and competition in a population (38, 39).

For accessory genes, ∼58% (n = 354/609) were found to have an essentiality score of 0 (Q1 = 0.0; median = 0.0; Q3 = 0.13). Only a single accessory gene was found in the highest quarter of the essentiality score range, compared to 310 core genes. This is consistent with the evolutionary conservation of core genes in the *S. pyogenes* population and suggests that loss of function of any of the 310 core genes is lethal for the organism. The single accessory gene found in the upper quarter of the essentiality score range was of particular interest as it is missing in some *S. pyogenes lineages* but is seemingly essential for growth in all strains and conditions tested in the transposon mutagenesis studies to date. The gene was annotated by EggNOG as *<u>s</u>ecretion and <u>a</u>cid tolerance D* (*satD*) (SF370 tag: SPY_RS04905/SPy1171). *satD* was found to be highly essential across *emm*1, *emm*28 and *emm*49 strains, but is an accessory gene across the population of complete *S. pyogenes* genomes (178/249, ∼71%). In *Streptococcus mutans*, *satD* and remaining *sat* genes have been associated with acid stress tolerance (40, 41). However, *satD* and related genes have not been studied in other streptococci. The example of *satD* showcases the importance of transposon sequencing meta-analyses, and how integration of a pangenomic framework can expose strain dependant adaptations otherwise missed when examining only a single strain or lineage in isolation.

### Stricter essentiality requirements under in vivo conditions

To assess gene essentiality between *in vivo* and *in vitro* conditions, we calculated the mean essentiality score within each condition irrespective of strain genotype. When comparing the distributions of essentiality of genes across *in vivo* (Q1 = 0.25; median = 0.25; Q3 = 1.25) and *in vitro* (Q1 = 0.00; median = 0.00; Q3 = 0.50) conditions, genes under *in vivo* conditions had a significantly higher essentiality score (paired Wilcoxon signed-rank test: p<2.2·10^-16^) (Figure 2A). The median essentiality was found to be 0.25 *in vivo* and 0.00 *in vitro*. These findings suggest that under *in vivo* conditions, *S. pyogenes* is more sensitive to gene knock-out and loss of function compared to *in vitro* conditions. Examining genes identified 245 (245/1,999, 12.3%) with large variation in essentiality score (>0.5) between *in vivo* and *in vitro* conditions (Figure 2B). Only three genes were of higher essentiality under *in vitro* conditions. This was contrasted with 242 genes with greatly increased essentiality score under *in vivo* conditions (Figure 2B, Supplementary Table 2, Supplementary html 2). Among the 242 genes, with greater essentiality *in vivo*, cellular processing, and signalling-related genes were overrepresented compared to metabolism (post-hoc Chi-squared: adjusted p=0.046), and information storage and processing (post-hic Chi-squared: adjusted p=0.007) related genes (Figure 2B). Using this metric we can hone into additional examples of poorly annotated core genes that may have important functions under *in vivo* conditions. One such example is a homolog to an RNA binding protein CvfD from *S. pneumoniae*, which can be found in *S. pyogenes* (SF370 tag: SPY_RS06755/SPy1619) (42). This gene is highly essential *in vivo* and moderately essential *in vitro*. In *S. pneumoniae*, *cvfD* has been linked to virulence in a mouse model of invasive pneumonia and is found to regulate 144 mRNA transcripts. The *S. pyogenes* homolog of *cvfD* is uncharacterized but could be important in regulating expression at a post-transcriptional level under *in vivo* conditions.

**Figure 2.**
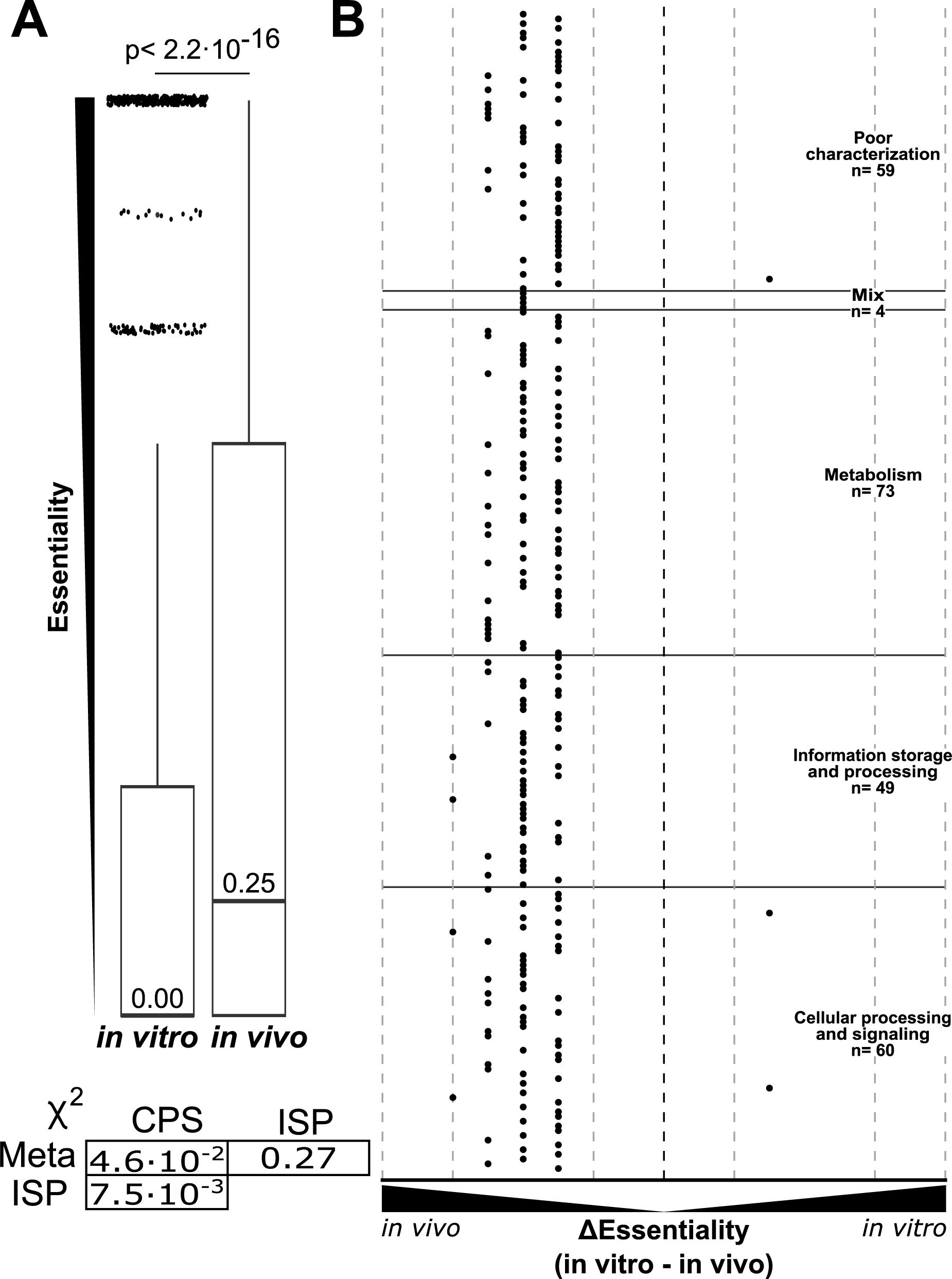
Differential gene essentiality across macro conditions. **A**: Boxplot of gene essentiality scores based on in vitro and in vivo experimental conditions. Outliers (essentiality score > 1.5 inner quartile range) are marked by black points. P-value for a Wilcoxon’s paired signed rank test between the two categories is given. **B**: distribution of genes with large difference in essentiality score (≥0.5) across in vitro and in vivo conditions (n = 245). Vertical dashed lines indicate differences of 2, 1.5, and 0.5, with the black dashed line indicating 0 difference between in vivo and in vitro. Major functional categories are given for points and separated by horizontal black lines. Points left of the black vertical dashed line have higher essentiality score in vivo. Conversely, points right of the line have increased essentiality score in vitro. In the lower left corner adjusted p-values for a Dunn’s test among well represent functional categories is given. Categories are shortened as follows: Metabolism (Meta), Information storage and processing (ISP), and Cellular processing and signalling (CPS).

### Contextualising gene essentiality with operon architecture

Many genes in bacteria are components of, and are evolutionary maintained within, polycistronic operons. These genomic features can be highly important due to transcriptional suppression and promotion mechanics and genome compactness. To assess operons and their essentiality, we used operon structures as defined in *emm*1 strain S119 (33). Of the 750 operons identified in S119, 726 (96.8%) were successfully mapped to the pangenome of the 249 complete genomes. Mapped S119 operons encompassed 405 monocistronic and 321 polycistronic operons. The systematic comparison of pangenomes and gene essentiality enables an interrogation of maintenance of essential genes in operon structures. The essentiality score of an operon was defined as the mean essentiality score for all genes in the operon. There was a wide range of operon essentiality scores across operons of differing sizes. Of monocistronic operons 15% were essential (essentiality score >1.5 n = 61/405) with 17% of polycistronic operons deemed essential (essentiality score >1.5 n = 56/321). Operon size and essentiality score did not seemingly correlate (Figure 3A). A linear regression model found the size of an operon to be a poor predictor of essentiality, R^2^ = 9.71•10^-3^. With multiple essential genes (n = 61) found in monocistronic operons and operon size found to be a weak predicter of operon essentiality, operon structure does not appear to be correlated with essentiality.

**Figure 3.**
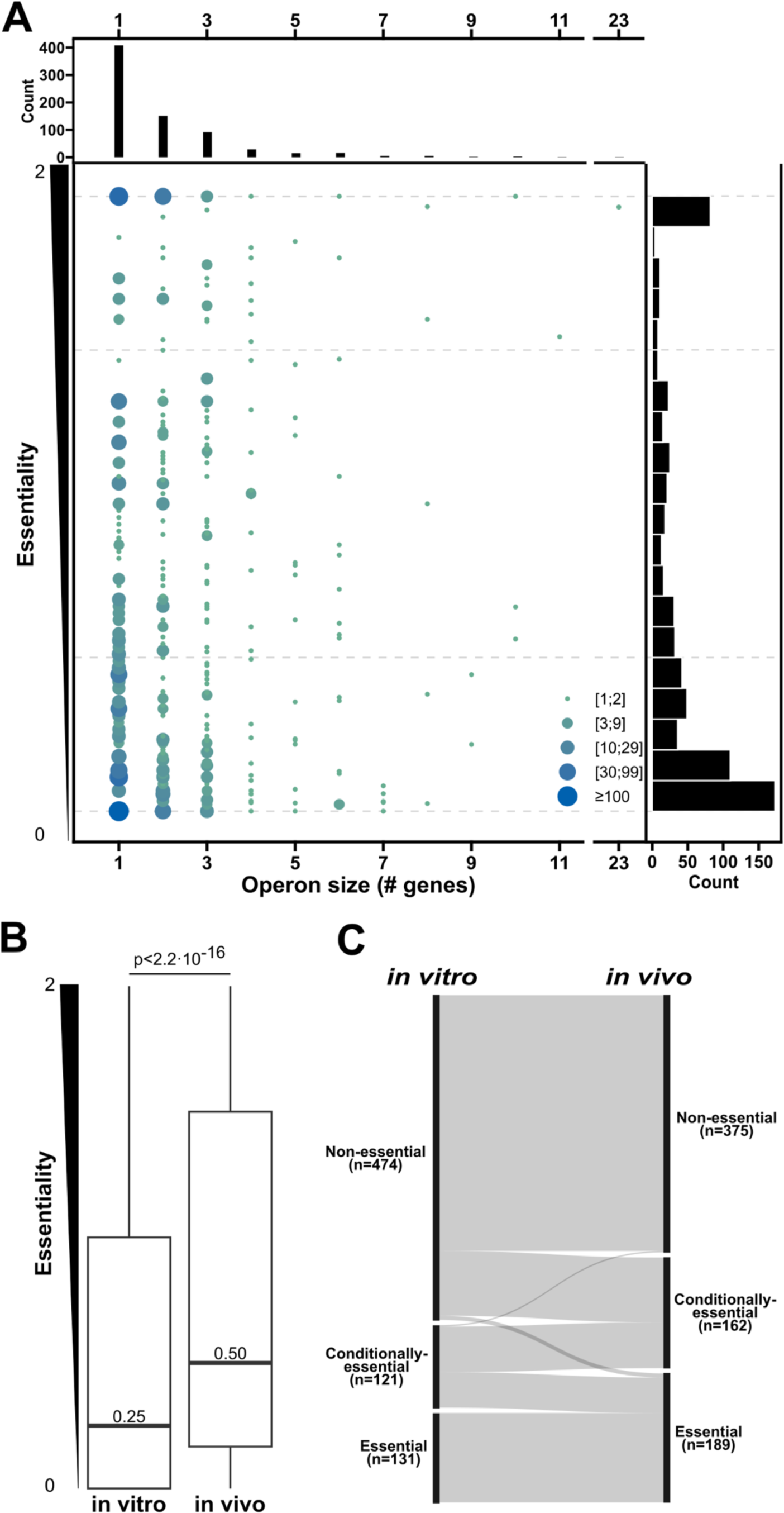
Operon essentiality for Streptococcus pyogenes. **A**: Distribution of size and essentiality score for operons (n = 732) from S119. Size and colour of points correspond to number of operons at a given position, relative to legend. Distributions for each variable is given along the outer axis. **B**: Boxplots for the distribution of essentiality score for operons under in vitro and in vivo conditions. Medians are given and the p-value for a paired Wilcoxon signed rank test is given. **C**: Sankey flow diagram for change of qualitative categories of essentiality scores for operons under in vitro and in vivo conditions. Categories are based on essentiality in the following order: Non-essential (essentiality score < 0.5), Conditionally-essential (0.5 ≤ essentiality score ≥ 1.5), Essential (essentiality score > 1.5). Grey links between the two columns (in vitro and in vivo) indicate operons position in the given essentiality categories across conditions.

We next examined the influence of growth conditions, we calculated an *in vivo* and *in vitro* essentiality score for each operon. Similar to findings for individual genes, comparing operon essentiality score under different conditions showed the *in vivo* essentiality score (Q1 = 1.67; median = 0.50; Q3 = 1.50) to be significantly higher than the *in vitro* score (Q1 = 0.25; median = 0.25; Q3 = 1.00) (paired Wilcoxon signed-rank test: p < 2.2·10^-16^) (Figure 3B).

Operons were next categorised into 3 qualitative groups based on essentiality score. Non-essential, operons with low essentiality score; Conditionally-essential, operons essential in some lineages but not all or resulting in decreased fitness and; Essential, operons with a high essentiality score across all strains. Division of operons into essentiality categories under *in vivo* and *in vitro* conditions separately demonstrated that operon essentiality can vary across conditions, with some operons being in different essentiality categories across *in vivo* and *in vitro* conditions (Figure 3C). These operons and the genes within them may be of interest in understanding the physiology and survival of *S. pyogenes*. As an example, the operon for the Arginine deaminase pathway (o_0507 in S119), which has previously described as important during skin infection (43), has an increase in essentiality score of ≥0.5 for all genes *in vivo* compared to *in vitro*, with an increase in essentiality ∼1 for the entire operon. Operons that have not been previously identified as important under *in vivo* conditions can also be discovered in the same way, potentially allowing new insights into *S. pyogenes* during infection conditions. One such is the S119 operon o_0575 encoding *pptA* and *pptB* (also called *ecsAB*, SF370 tag: SPY_RS07195/SPy1728, and SPY_RS07200/SPy1729). These two genes have been shown to export competence pheromones in *S. pyogenes* (44) but have not been linked to importance under *in vivo* conditions. *pptAB* are of low essentiality under *in vitro* conditions but increase to a moderate essentiality score under *in vivo* conditions. In *Streptococcus equi* subspecies *equi pptAB* have been shown to be important for survival in horse blood cultures but not in hydrogen peroxide (45).

## Discussion

One of the challenges of bacterial population biology is to identify important common pathways among populations where gene content is variable, and evolution is not static. To facilitate broader analytical resources to the *S. pyogenes* research community, systematic analytical frameworks of functional genomic studies such as transposon mutagenesis studies undertaken across different conditions and strain backgrounds represent an important tool in defining gene essentiality in *S. pyogenes*. This re-analysis identified significant differences between essentiality scores of core genes compared to accessory genes. The identification of a single highly essential accessory gene, *satD*, highlights how the framework presented here, using a pangenome and transposon sequencing, allow for investigation of possible lineage adaptations encoded by accessory genes. Further, integration of operon structures of *S. pyogenes* indicated different requirements between *in vivo* and *in vitro* conditions, with operons having a significantly greater essentiality score under *in vivo* conditions.

The fact that *S. pyogenes* is divided into hundreds of genomic lineages with varying and often poorly characterised gene content complicates the effort to fully understand this human pathogen (2). This emphasises the epidemiological differences between *S. pyogenes* lineages that have been puzzling researchers for decades (46, 47). However, with the sparse number of datasets (n = 9) and *emm* lineages (n = 3) represented in transposon sequencing studies, much is still to be learned about lineage-specific differences in essentiality of genes in the context of adaptation to different conditions or niches. In this work, only cross lineage, *in vivo,* and *in vitro* essentiality scores were calculated. With future studies, inclusion of more strains and conditions would provide new opportunities to interpret essentiality across lineages. Additionally, essentiality scores within the same strain compared across *in vivo* and *in vitro* conditions could give a deeper understanding of how a strain adapts to these two different macro conditions.

Little work has been done to investigate how batch effects may influence transposon sequencing, and how to control them. The Zero-Inflated Negative Binomial method from the Transit package, used to analyse Himar1 like transposon libraries, allows for co-variates to be included in the statistical model, but this requires a careful experimental design and does not lend itself to this meta-analysis (28). Other biases known to affect transposon sequencing studies are the choice of transposon and the subsequent method of analysing the differential presence of transposons across the genome (13, 28). How to best handle specific transposons of choice is an ongoing field of research (13, 48). In this paper we have chosen methods specifically developed to analyse transposon sequencing data, compared to the common but likely problematic approach of using RNA-seq analysis methods. The methods used here incorporate information on the transposon used in producing the data and hence reduce the bias that can be introduced from the method of analysis.

Here we have undertaken a major reanalysis and assessed results from a repository of gene essentiality across *emm* lineages and conditions for *S. pyogenes*. This has been integrated into a pangenome of complete genomes and operon structure to increase the usefulness of the dataset. Additionally, the method used here to calculate essentiality can be extended to include new studies and update essentiality scores. Future studies will hopefully broaden our understanding on mechanisms underlying gene essentiality across additional strains and conditions, as have been done in other species (24–26).

## Materials and methods

### Pangenome construction and functional annotation

255 complete genomes of *Streptococcus pyogenes* were collected from RefSeq as of 15th Janurary 2021 year. Quality control of misassembles was done using Socru v.2.2.4 (49) and by identifying spurious additional chromosomes (contigs) not related to extra chromosomal elements meant that the final number of reference genomes analysed in this study was 249. A pangenome was constructed using Panaroo and flags: clean-mode set to strict, initial percent identity to 90% (-i 0.9), and maximum length difference to 95% (-l 0.95) (29). Functional annotations for each pangenome cluster identified by Panaroo were obtained by running EggNOG-mapper v.2.1.3 using the EggNOG database v5.0 (30, 31).

### Transposon sequencing analysis

Raw sequencing data was acquired from the sequence read archive (SRA) using the SRA Toolkit (github.com/ncbi/sra-tools). FastQC v.0.11.9 (www.bioinformatics.babraham.ac.uk/projects/fastqc/) and seqkit v.2.2.0 were used to assess quality of raw fastq files (50). Depending on transposon type (Tn5 or Himar1) and study, data were trimmed to isolate genomic DNA from reads using Cutadapt v.3.8.6 (51) [https://journal.embnet.org/index.php/embnetjournal/article/view/200]. Tn5 like transposons were trimmed using the sequence AGATCGGAAGAG, and flags: -e 2, --no-indels, -O 10, and --action trim. For Untrimmed Himar1 transposon reads the sequence: ACAGGTTGGATGATAAGTCCCCGGTCTGACACATCTCCCTAT was used with flag: -m 10. Subsequent quality filtering was conducted using seqkit seq -t dna -Q 34. Alignment and counting of reads were done using the Transit TPP tool v.3.2.6 modified to run bowtie v.1.3.1 as the alignment tool, using flags: -S, -m 1, -k 1, -n 0 (27, 28, 52). For each lineage used in the original studies a corresponding RefSeq reference genome was used for alignment and construction of Prot_tables (Transit specific annotations format) related to the pangenome annotations (NZ131: GCF_000018125.1, GCF_, Alab49: GCF_000230295.1, 5448: GCF_001021955.1, MGAS2221: GCF_012572265.1, MGAS27961: GCF_004010855.1). For all input conditions of Tn5 libraries essential genes were determined using the Tn5Gaps (Tn5 methods from Transit (53)). To identify differential abundance of Tn5 insertions across input and output libraries the Resampling method from Transit was used (27). For Himar1 libraries essential genes and differences between input and output libraries were determined using the Zero-Inflated Negative Binomial method (ZINB) from Transit (54). All datasets were normalized using the beta geometric distribution (-norm betageom).

### Analysis of gene essentiality

Results from methods identifying essential and conditionally-essential genes were processed in R v.4.1.1 and RStudio v.1.4.17. Genes in the ZINB output of Transit marked as ‘pan-essential’ or ‘pan-growth defect’ in the status columns were classified as essential. Genes that did not meet this criterion but had adjusted p-values < 0.05 and log2(fold change) ≤ 0 were marked as conditionally-essential.

The FactorMineR v.2.8 package in R was used to conduct the quality assurance using multiple correspondence analysis (55). Essentiality calls were integrated using information from the pangenome and combined with S119 operons structures (33). To calculate essentiality scores for genes across lineages and conditions the essentiality calls from transposon sequencing results were used. The three groups, non-essential, conditionally-essential, and essential, were seen as ordered (ordinal categorical data). Categories were assigned numeric values based on their order (non-essential = 0, conditionally-essential = 1, essential = 2). Using the numeric values of essentiality, the lineage weighted essentiality score was calculated in a two-step process. First, for each gene the mean essentiality score within lineages (*emm*1, *emm*28 or *emm*49) was calculated. Secondly, the essentiality score across lineages was calculated as the mean of all lineages means for the given gene. For *in vivo* and *in vitro* essentiality scores the mean for all datasets belonging to either one of the two was calculated for each gene, without correction for lineage. Having calculated an essentiality score, genes were defined as essential if an essentiality score ≥1.5. An essentiality score < 1.5 and > 0.5 were considered as conditionally-essential, and essentiality score ≤ 0.5 was considered non-essential.

### Plotting, data wrangling and statistical tests

Plots were produced using ggplot2 v.3.4.2, ggprism v.1.0.4, ggbreak v.0.1.2, networkD3 v.0.4, and htmlwidgets v.1.6.2 in R (56–60). Data handling was aided by the R package: reshape2 v.1.4.4 (61). Interactive html tables were produced using the DT v.0.28 package in R (62). Statistical tests were conducted in R. Wilcoxon test to compare medians of two samples were carried out using wilcox.test(), as a paired test (paired=T) if specified.

Test for difference in median essentiality score across major COG categories was conducted using a Kruskal-Wallis text using kruskal.test(). In case of a significant (p<0.05) Kruskal-Wallis test, a Dunn’s test was used as post-hoc analysis with Benjamini-Hochberg p-value correction (dunn.test() from the dunn.test R package v.1.3.5)(63). Test for overrepresentation of major COG categories was conducted using Chi-squared tests, with no correction (correct = F). Upon a significant comparison across multiple groups (p<0.05) individual Chi-squared comparisons between categories were made with subsequent Benjamini-Hochberg p-value correction. Linear regression for operon size and essentiality association was conducted using the lm() function in R, with operon essentiality as a function of operon size without fixing the intercept of the regression to the origin.

## Data availability statement

Data on the essentiality of genes and operons are provided as interactive html files as well as comma separated tables. An additional table relating the pangenome cluster to locus_tags of reference genomes used is provided. Genes annotated with no gene name are in the pangenome given a name of group_####. Therefore, searching directly for a given gene name may not be possible.

## Supporting information

Operon essentiality table

Pangenome essentiality table

Pangenome relation to references

Operon essentiality table - interactive

Pangenome essentiality table - interactive

Pangenome relation to references - interactive

## Acknowledgement

The authors would like to thank Ouli Xie for critical reading and feedback on the manuscript. MGJ is supported by The Melbourne Research Scholarship from The University of Melbourne. MRD was supported by a University of Melbourne CR Roper Fellowship. This research received no specific grant from any funding agency in the public, commercial, or not-for-profit sectors.

---

**Supplementary figure 1.**
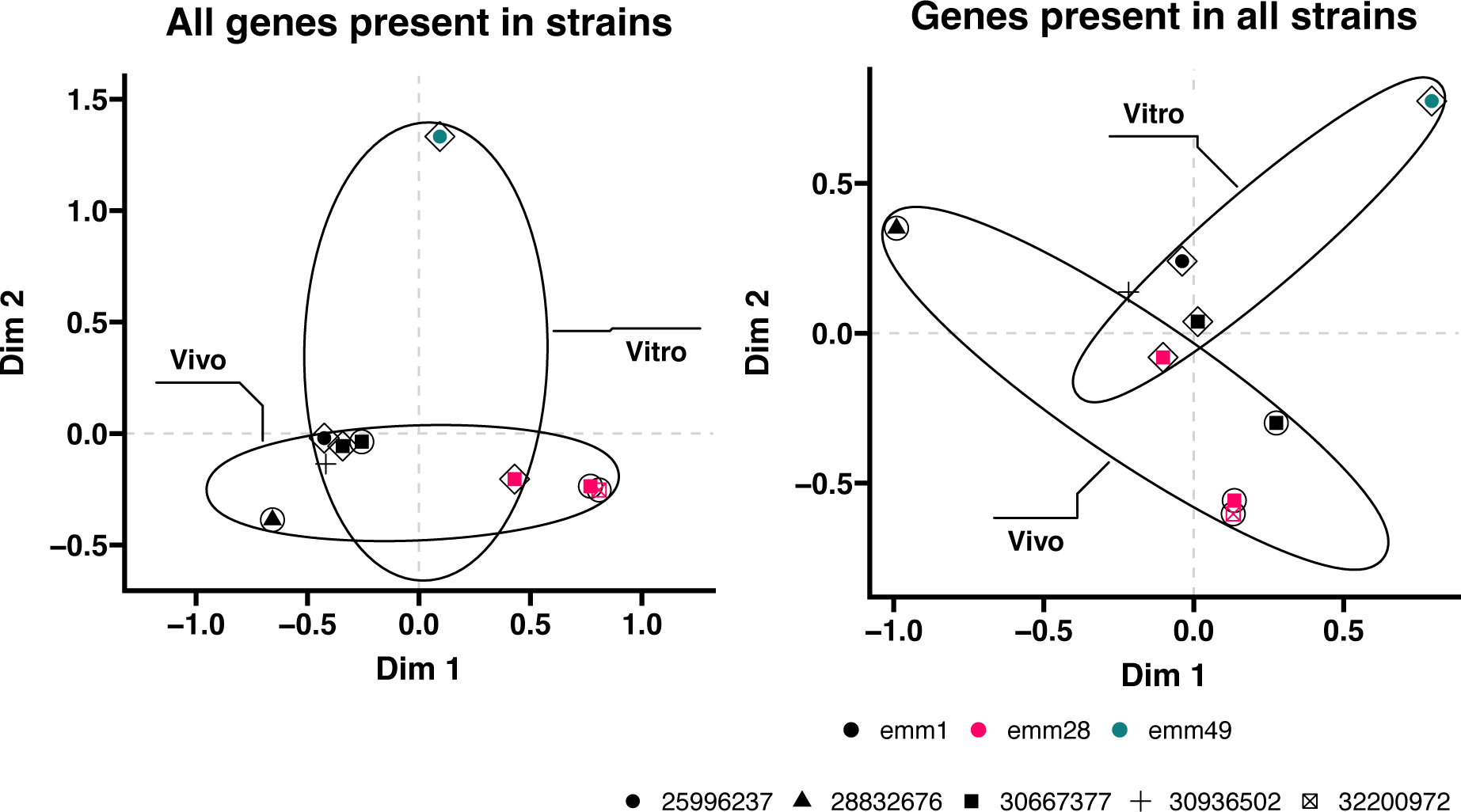
Multiple correspondence analysis of transposon sequencing data. Each plot illustrates the first and second dimension returned from a multiple correspondence analysis of essentiality group for all genes present in strains (left) and genes present in all strains (right). Ellipses surround samples from in vivo and in vitro conditions. Point colours indicate emm lineage used in a dataset, and point shape indicate the PubMed ID from which data was obtained.

